# Lipid Alterations in African American Prostate Cancer

**DOI:** 10.1101/2021.11.05.467481

**Authors:** Anindita Ravindran, Danthasinghe Waduge Badrajee Piyarathna, Jie Gohlke, Vasanta Putluri, Tanu Soni, Stacy Lloyd, Patricia Castro, Subramaniam Pennathur, Jeffrey A Jones, Michael Ittmann, Nagireddy Putluri, George Michailidis, Thekkelnaycke M Rajendiran, Arun Sreekumar

**Author notes:** **Correspondence** Arun Sreekumar, Ph.D., Charles C Bell Endowed Professor, Center for Metabolism and Experimental Therapeutics, Department of Molecular and Cell Biology, Baylor College of Medicine, Rm 120 DJ, Jewish Biomedical Research Building, One Baylor Plaza, Houston, TX-77030. Equal First Authors (Anindita Ravindran, Danthasinghe Waduge Badrajee Piyarathna, Jie Gohlke). Equal Senior Authors (George Michailidis, Thekkelnaycke M Rajendiran, Arun Sreekumar). **Author Contributions** A.S.K and G.M. designed the study. A.S.K. supervised the study. J.G. and V.P. performed the mass spectrometry experiments. S.T, T.M.R, S.P and N.P. helped develop the lipidomics platform and associated data analysis pipeline, D.W.B.P. and S.L. analyzed the data, M.I., P.C., and J.A.J contributed clinical samples and clinical insights; A.R., S.L., J.G., J.A.J., S.P., S.L., G.M., A.S.K. provided inputs for data interpretation; D.W.B.P. and A.R. generated the figures; A.R., J.G., G.M., and A.S.K. wrote the paper.

## Abstract

African-American (AA) men are more than twice as likely to die of prostate cancer (PCa) than European American (EA) men. Previous *in-silico* analysis revealed enrichment of altered lipid metabolic pathways in pan-cancer AA tumors. Here, we performed global unbiased lipidomics profiling on 48 matched localized PCa and benign adjacent tissues (30 AA, 24 ancestry-verified, and 18 EA, 8 ancestry verified) and quantified 429 lipids belonging to 15 lipid classes. Significant alterations in long chain polyunsaturated lipids was observed between PCa and benign adjacent tissues, low and high Gleason tumors, as well as associated with early biochemical recurrence, both in the entire cohort, and within AA patients. Altered levels of cholesteryl esters, and phosphatidyl inositols delineated AA and EA PCa, while levels of triglycerides, phosphatidyl glycerol, phosphatidyl choline, phosphatidic acid and cholesteryl esters distinguished AA and EA PCa patients with biochemical recurrence. These first-in-field results implicate lipid alterations as biological factors for prostate cancer disparities.

## Introduction

Prostate cancer (PCa) progression is twice as aggressive and more than twice as morbid in African Americans (AA) compared to European Americans (EA), yet the specific biochemical basis underlying this disparity remains unknown^1^. While environmental factors such as diet and access to proper healthcare may be significant causes for increased mortality rates associated with PCa in AA men, recent reports suggest that metabolic alterations may also drive disparity in PCa progression^2^. Our previous *in silico* study described enriched adipogenesis, fatty acid metabolism and cholesterol homeostasis, among others, as being enriched in AA tumors across multiple different tumor types, which was further validated in an independent PCa gene expression dataset^3^. In addition, elevated cholesterol levels in serum have been reported to be associated with increased biochemical recurrence (BCR) specifically in AA men with PCa^4^. Here, for the first time we describe race-specific clinically relevant alterations in PCa lipidome.

## Results

We performed unbiased global lipidomic profiling using Triple Time Of Flight (Triple-TOF) liquid chromatography-mass spectrometry (LC-MS) on 48 localized PCa and matched benign adjacent tissue (**Table 1** for Clinical Information, 30AA and 18 EA pairs, 24 AA and 8 EA tissues were ancestry verified). The profiling platform was highly reproducible with a % CV of 2.7 % in the liver pool (n= 24) for 429 lipids measured, CV of 5.3% for internal standards (n=26, **Supplementary Figure 1**) and normal distribution of the lipids overall and within each class (**Supplementary Figure 2**). The 429 lipid species identified belonged to 15 lipid classes including cholesteryl esters (CE), diacylglycerols (DG), lysophosphatidyl cholines (L-PC), lysophosphatidyl ethanolamines (L-PE), phosphatidic acid (PA), phosphatidyl cholines (PC), phosphatidyl ethanolamines (PE), phosphatidyl glycerol (PG), phosphatidyl inositols (PI), plasmenyl-phosphatidyl cholines (P-PC), plasmenyl-phosphatidyl ethanolamines (P-PE), phosphatidyl serines (PS), sphingomyelins (SM), and triglycerides (TG) (**Supplementary Tables 1-3**).

**Table 1.** Clinical parameters of prostate tumor samples used for lipidomics profiling. *ŷ represents the estimated value of the response variable.

| Variable | African American<br>(n=30) | European American<br>(n=18) |
| --- | --- | --- |
| Gleason Grade | Low ($\leq 6$): 20 | Low ($\leq 6$): 6 |
| | High ($\geq 7$): 10 | High ($\geq 7$): 12 |
| Recurrence | 5 | 5 |
| No Recurrence | 10 | 5 |
| West African, $\hat{y}^*$ | $0.8 \pm 0.1$ | $0.02 \pm 0.02$ |
| European, $\hat{y}^*$ | $0.2 \pm 0.1$ | $0.02 \pm 0.02$ |
| Native American, $\hat{y}$ | $0.0 \pm 0.0$ | $0.95 \pm 0.03$ |
| Genetic Ancestry<br>Verified (n) | 24 | 8 |

Principal Component Analysis (PCA) using all the lipids revealed a moderate separation of the benign and cancer samples, both across the entire cohort and within AA patients, suggestive of alterations in lipid components between these two tissue types (**Figure 1A and Supplementary Figure 3A**). Similarly, across the entire cohort as well as with AA patients, comparing PCa with benign adjacent tissue, the significantly altered lipids (FDR <0.1) within each lipid class were predominantly polyunsaturated (>2 double bonds) with long chain fatty acid chains (20-40 fatty acids, **Figures 1B, C** and **Supplementary Figures 3B, C, 4 and 5**) for altered lipids with FDR < 0.25). Included here were prominent elevation in levels of CE, TG and reduced levels of SM, PS, P-PE, PI and PG (**Figure 1C** and **Supplementary Figure 3C, Supplementary Figures 4 and 5, Supplementary Tables 1 and 2**). In addition, reduced levels of L-PC (across all PCa and within AA PCa), and SM (only within AA PCa), and significantly elevated levels of TG (only within AA PCa) were significantly associated with high Gleason tumors (Gleason >7 or =4+3, **Figure 1D** and **Supplementary Figure 3D**, all FDR<0.25). Further, across all PCa, lower levels of L-PC, SM, and PE were significantly associated with early biochemical recurrence (BCr, 5 years post-prostatectomy, FDR<0.35), a clinical indicator of aggressive PCa (**Supplementary Figure 3D**). Similarly, reduced levels of L-PC and P-PC, and elevated levels of PI were associated with early BCr with AA PCa patients (FDR<0.35, **Figure 1E**).

**Figure 1.**
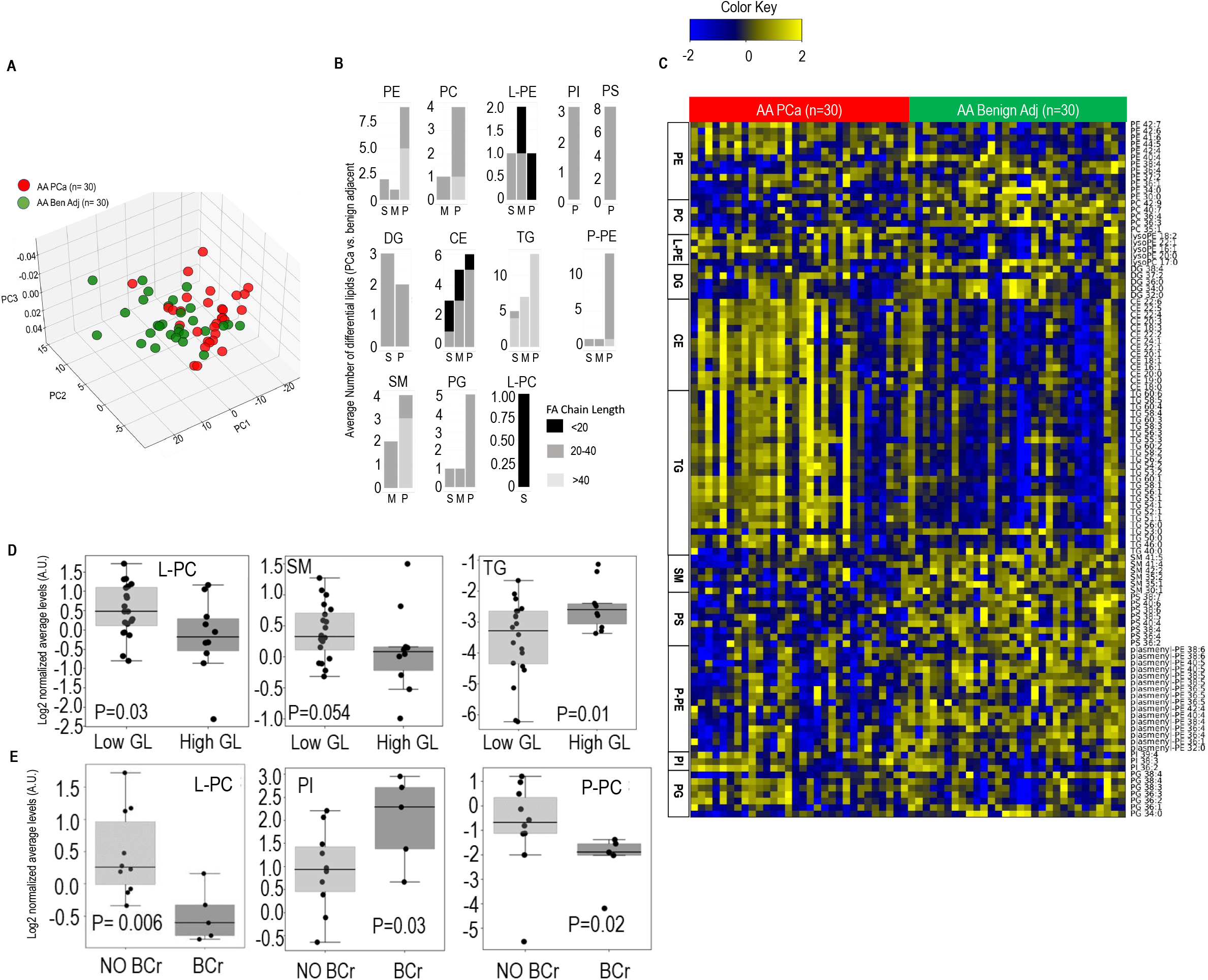
Altered lipidome in AA PCa vs matched benign patient prostate tissue. **(A)** PCA plot using lipid profiles in 30 paired AA PCa and benign adjacent prostate tissues. **(B)** Number of altered lipids within each class stratified by fatty acid chain length (see legend) and degree of saturation. S: Saturated, M: Mono-unsaturated, P: Poly-unsaturated (≥ 2 double bonds). **(C)** Heat map showing significantly altered lipids in AA PCa vs matched benign patient prostate tissues. Shades of Yellow and Blue represent up and down regulated lipids, respectively (see color key). Lipids are arranged by classes. PE: Phosphatidyl ethanolamine, PC: Phosphatidyl choline, PI: Phosphatidyl inositol, PG: Phosphatidyl Glycerol, PS: Phosphatidyl Serine, L-PE: Lyso Phosphatidyl Ethanolamine, TG: Triglycerides, SM: Sphingomyelin, P-PE: Plasmenyl Phosphatidyl Ethanolamine, CE: Cholesteryl Esters, DG: Diglycerides, L-PC: Lyso phosphatidyl choline. **(D)** Levels of L-PC (p=0.03) and SM (p=0.054) are significantly down-regulated, and levels of TGs (p=0.01) are significantly elevated in high Gleason grade (High GL, n=10) compared to Low Gleason grade (Low GL, n=20) tumors. **(E)** Lower levels of L-PC (p=0.006) and PL-PC (p=0.02) and elevated levels of PI (p=0.03) are associated with biochemical recurrence (BCr, 5 years post-prostatectomy) in PCa patients. No BCr (n=10); BCr (n=5). For panels B and C, paired t-test followed by Benjamini Hochberg (BH) False Discovery Rate (FDR<0.1) correction was used to compute differential analysis. For panels D and E, Mann-Whitney test with BH FDR<0.25 and <0.35 respectively were used to compute statistical significance.

Intriguingly, there were key changes in lipid profiles comparing Gleason grade matched AA (n=30) and EA (n=18) tumors. As shown in **Figure 2A**, PCA analysis of the lipid profiles was able to delineate AA and EA tumors for the most part. Furthermore, lipids (predominantly polyunsaturated and long chain) belonging to 10 lipid classes significantly distinguished AA and EA PCa (**Figure 2B, Supplementary Figure 6A**), that included prominent elevation (FDR < 0.1) in levels of CE, TG, PI and PA class of lipids (**Figure 2C**, altered lipids with FDR<0.25 in **Supplementary Figure 6, Supplementary Table 3**). Interestingly, comparison of AA and EA PCa patients with early BCr revealed conspicuous elevation (FDR<0.35, altered at least two-fold) in levels of TG and CE, in the former, suggesting race-specific de-regulation of different lipid pathways in aggressive tumors (**Figure 2D**)..

**Figure 2.**
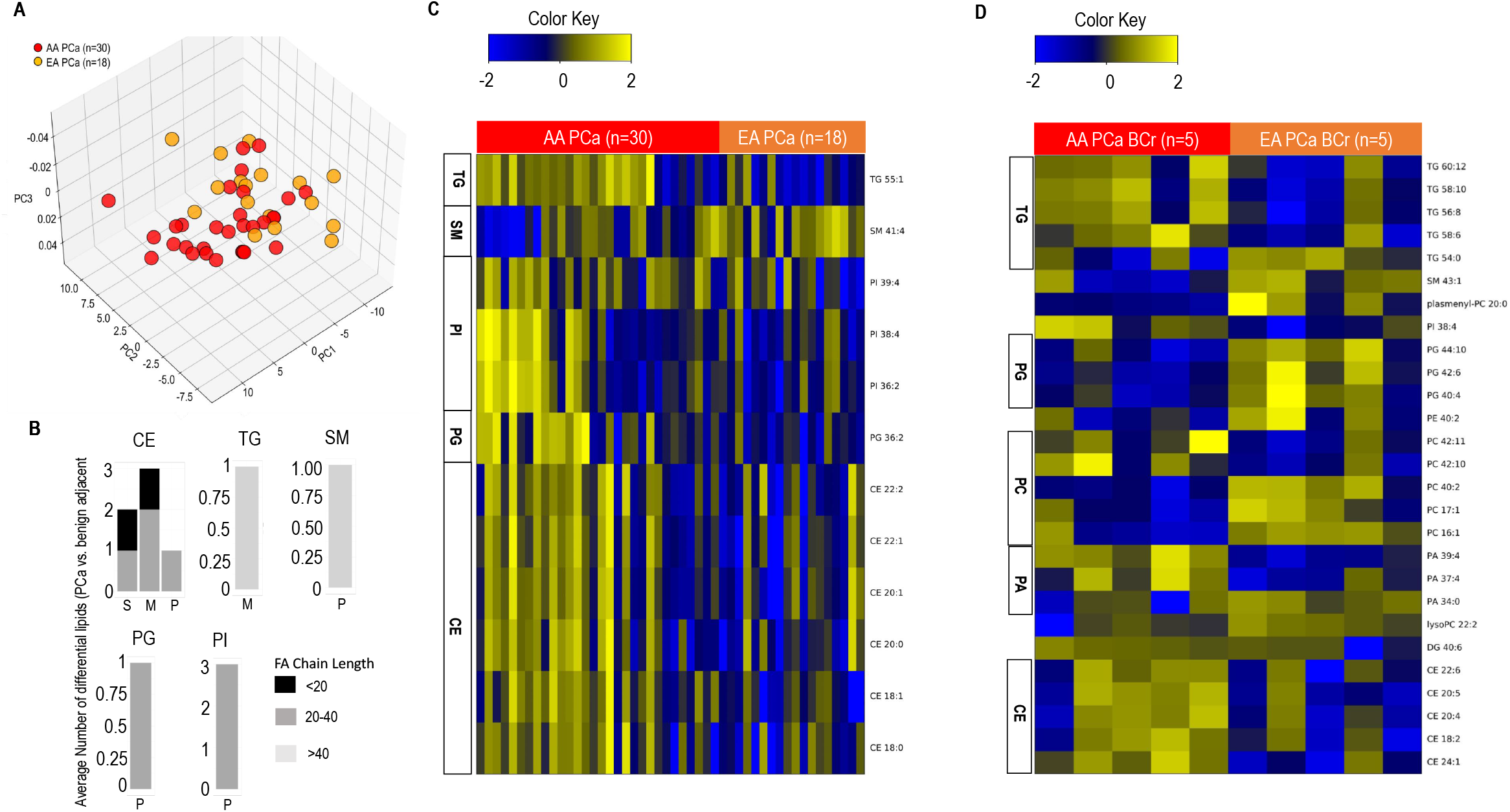
Altered lipidome in AA PCa vs EA PCa patient prostate cancer tissue. **(A)** PCA plot using lipid profiles in 30 AA and 18 EA PCa tissues **(B)** Number of altered lipids within each class stratified by fatty acid chain length (see legend) and degree of saturation. S: Saturated, M: Mono-unsaturated, P: Poly-unsaturated (≥ 2 double bonds). **(C)** Heat map showing significantly altered lipids in AA PCa vs EA PCa tissues. Shades of Yellow and Blue represent up and down regulated lipids, respectively (see color key). **(D)** Heat map showing differential lipids comparing AA PCa and EA PCa with biochemical recurrence within 5 years post-prostatectomy. Lipids are arranged by classes. P-PC: Plasmenyl Phosphatidyl choline, P-PE: Plasmenyl Phosphatidyl Ethanolamine, TG: Triglycerides, SM: Sphingomyelin, PI: Phosphatidyl inositol, PG: Phosphatidyl Glycerol, PC: Phosphatidyl choline, CE: Cholesteryl Esters, PA: Phosphatidic Acid. For panels B and C, paired t-test coupled to Benjamin Hochberg False Discovery Rate (FDR<0.1) correction was used to compute differential analysis. For panel D, Mann-Whitney test coupled to Benjamin Hochberg False Discovery Rate (FDR<0.35) correction was used to compute differential analysis. Lipids that were altered at least 2-fold with a FDR corrected p-value of P<0.05 are shown.

## Discussion

Our finding describes unique race-associated alterations in specific lipid classes that include increase in CEs, PI, and TGs in PCa. Interestingly, both CE and TGs were prominently higher in AA vs EA patients with early BCr.

CEs are the primary source of cholesterol for steroidogenesis^5^. CEs (cholesterol oleate in particular) have been detected in a previous lipidomic screen and proposed as potential molecular biomarkers for PCa^6^. Increases in CEs were also shown to be associated with PCa tumor progression and metastasis and aberrant accumulation of CEs was found to be correlated with PTEN loss and PI3K/Akt activation^7,8^. This activation of the PI3K/Akt pathway, which has recently been demonstrated to be higher in AA PCa, is corroborated by elevated PI (an important second messenger in the PI3K/Akt pathway) levels in these tumors in our analysis. Furthermore, this is consistent with our previous publication where we reported that the oncogene MNX1 is upregulated in AA men in response to activated PI3K/AKT signaling and drives lipid synthesis via activation of SREBP1 and fatty acid synthase^9^.

In addition, high levels of TGs seen in AA PCa are thought to be associated with PCa progression^10^. TGs are an important energy source when hydrolyzed into free fatty acids (FFAs). Cancer cells use the energy generated from these TG pools to drive tumor proliferation, specifically under hypoxia^11^.

Intriguingly, increase in CE and TG in AA vs EA patients with early BCr is a novel finding that could be regulated by androgens that are known to affect triglyceride and cholesteryl ester pools by contributing to lipid synthesis^12^.

Interestingly, in our study, the majority of the altered lipids in the context of race in PCa include polyunsaturated fatty acids (PUFAs). PUFAs are known to be high in AA diet which predominantly comprises of corn and soy^13^. Moreover, AA men are known to have significantly higher circulating levels of arachidonic acid (a long chain PUFA implicated in inflammation) than EA men^14^. Our finding that the AA PCa lipidome is rich in long chain PUFAs further supports these prior observations and potentially reveals an impact of diet on PCa risk or progression.

The high incidence of PCa as well as the aggressive nature of tumor progression in AA men make it imperative to seek novel biological means to comprehend the disparity. Understanding the alterations in lipid composition and thereby discovering lipid biomarkers that are prognostic for PCa progression in AA but not EA men may be critical to developing race-associated therapeutic strategies. Our finding of increased CEs and TGs in AA PCa strongly supports the use of these specific lipid classes as prognostic biomarkers for aggressive PCa progression.

Taken together, our descriptive data for the first time reveals key changes in PCa lipidome in AA compared to EA patients. Additional studies are needed to address these key findings in the context of PCa disparities.

## Methods

Lipid extraction from tissues and mass spectrometry analysis were performed as described previously^15–18^. Missing values were imputed using the K nearest-neighbor method (KNN). Data was log2 transformed and day median normalized and checked to ensure normality, both with in each lipid class and across all lipids measured (**Supplementary Figure 2**). Unless specified, differential lipids were determined using paired t-test (p<0.05), followed by the Benjamini-Hochberg (BH) procedure for false discovery rate (both FDR <0.1 and FDR<0.25 have been shown) correction. Lipid profiles comparing AA vs EA PCa was determined by subtracting their respective paired benign values, and using a permutation test (p <0.05, FDR<0.25). Mann-Whitney statistics with BH FDR correction was used to determine lipid classes that distinguished PCa by Gleason and BCr. FDR<0.25 and FDR<0.35, respectively were used to determine altered lipids in comparisons involving Gleason and BCr.

The normalized data for this study has been deposited to the Metabolomics Workbench (https://www.metabolomicsworkbench.org/) under the trackID 2901.

