## Supplementary Tables for "Lipid Alterations in African American Prostate Cancer"

**Supplementary Table 1. List of differential lipids (FDR<0.25) in PCa vs adjacent benign tissues.** List of detected differential lipids between PCa and matched benign adjacent tissue arranged by lipid name, length, and bond number. PE: Phosphatidyl ethanolamine, PC: Phosphatidyl choline, PI: Phosphatidyl inositol, PG: Phosphatidyl Glycerol, PS: Phosphatidyl Serine, L-PE: Lyso-Phosphatidyl Ethanolamine, TG: Triglycerides, SM: Sphingomyelin, P-PE: Plasmenyl-Phosphatidyl Ethanolamine, CE: Cholesteryl Esters, DG: Diglycerides, CL: Cardiolipins, L-PC: Lyso-Phosphatidyl Choline. The fatty acid chain length and the number of bonds (degree of saturation) are indicated.

| Lipid Name | Chain Length | Bond Number |
|---|---|---|
| CE 16:1 | 16 | 1 |
| CE 18:0 | 18 | 0 |
| CE 18:1 | 18 | 1 |
| CE 18:3 | 18 | 3 |
| CE 19:0 | 19 | 0 |
| CE 20:0 | 20 | 0 |
| CE 20:1 | 20 | 1 |
| CE 20:3 | 20 | 3 |
| CE 22:1 | 22 | 1 |
| CE 22:2 | 22 | 2 |
| CE 22:4 | 22 | 4 |
| CE 22:5 | 22 | 5 |
| CE 22:6 | 22 | 6 |
| CE 24:1 | 24 | 1 |
| DG 32:0 | 32 | 0 |
| DG 34:0 | 34 | 0 |
| DG 35:1 | 35 | 1 |
| DG 35:2 | 35 | 2 |
| DG 36:0 | 36 | 0 |
| DG 36:4 | 36 | 4 |
| DG 37:1 | 37 | 1 |
| DG 37:2 | 37 | 2 |
| DG 37:4 | 37 | 4 |
| DG 38:1 | 38 | 1 |
| DG 38:2 | 38 | 2 |
| DG 38:4 | 38 | 4 |
| DG 38:5 | 38 | 5 |
| DG 38:6 | 38 | 6 |
| DG 40:2 | 40 | 2 |
| DG 40:5 | 40 | 5 |
| lysoPC 17:0 | 17 | 0 |
| lysoPE 16:1 | 16 | 1 |
| lysoPE 18:2 | 18 | 2 |
| lysoPE 20:3 | 20 | 3 |
| lysoPE 22:1 | 22 | 1 |

| | | |
|---|---|---|
| lysoPE 22:6 | 22 | 6 |
| PA 37:4 | 37 | 4 |
| PC 28:0 | 28 | 0 |
| PC 31:0 | 31 | 0 |
| PC 32:0 | 32 | 0 |
| PC 33:0 | 33 | 0 |
| PC 34:1 | 34 | 1 |
| PC 34:4 | 34 | 4 |
| PC 35:1 | 35 | 1 |
| PC 36:0 | 36 | 0 |
| PC 36:3 | 36 | 3 |
| PC 36:4 | 36 | 4 |
| PC 37:5 | 37 | 5 |
| PC 38:1 | 38 | 1 |
| PC 38:4 | 38 | 4 |
| PC 38:8 | 38 | 8 |
| PC 40:10 | 40 | 10 |
| PC 40:4 | 40 | 4 |
| PC 40:6 | 40 | 6 |
| PC 40:7 | 40 | 7 |
| PC 40:8 | 40 | 8 |
| PC 42:9 | 42 | 9 |
| PE 30:0 | 30 | 0 |
| PE 32:2 | 32 | 2 |
| PE 33:2 | 33 | 2 |
| PE 34:0 | 34 | 0 |
| PE 35:3 | 35 | 3 |
| PE 36:1 | 36 | 1 |
| PE 36:3 | 36 | 3 |
| PE 36:4 | 36 | 4 |
| PE 37:2 | 37 | 2 |
| PE 37:4 | 37 | 4 |
| PE 38:4 | 38 | 4 |
| PE 38:5 | 38 | 5 |
| PE 39:4 | 39 | 4 |
| PE 40:1 | 40 | 1 |
| PE 40:3 | 40 | 3 |
| PE 40:4 | 40 | 4 |
| PE 41:4 | 41 | 4 |
| PE 41:6 | 41 | 6 |
| PE 42:4 | 42 | 4 |
| PE 42:6 | 42 | 6 |
| PE 42:7 | 42 | 7 |

| | | |
|---|---|---|
| PE 44:5 | 44 | 5 |
| PG 34:0 | 34 | 0 |
| PG 34:1 | 34 | 1 |
| PG 36:1 | 36 | 1 |
| PG 36:3 | 36 | 3 |
| PG 38:4 | 38 | 4 |
| PG 38:4 | 38 | 4 |
| PG 44:11 | 44 | 11 |
| PI 36:2 | 36 | 2 |
| PI 38:3 | 38 | 3 |
| PI 40:4 | 40 | 4 |
| plasmenyl-PC 18:0 | 18 | 0 |
| plasmenyl-PE 32:0 | 32 | 0 |
| plasmenyl-PE 36:1 | 36 | 1 |
| plasmenyl-PE 36:4 | 36 | 4 |
| plasmenyl-PE 36:4 | 36 | 4 |
| plasmenyl-PE 36:5 | 36 | 5 |
| plasmenyl-PE 36:5 | 36 | 5 |
| plasmenyl-PE 36:5 | 36 | 5 |
| plasmenyl-PE 38:4 | 38 | 4 |
| plasmenyl-PE 38:5 | 38 | 5 |
| plasmenyl-PE 38:5 | 38 | 5 |
| plasmenyl-PE 38:6 | 38 | 6 |
| plasmenyl-PE 38:6 | 38 | 6 |
| plasmenyl-PE 40:4 | 40 | 4 |
| plasmenyl-PE 40:5 | 40 | 5 |
| plasmenyl-PE 40:5 | 40 | 5 |
| plasmenyl-PE 42:4 | 42 | 4 |
| PS 36:2 | 36 | 2 |
| PS 36:4 | 36 | 4 |
| PS 38:4 | 38 | 4 |
| PS 38:5 | 38 | 5 |
| PS 38:6 | 38 | 6 |
| PS 38:7 | 38 | 7 |
| PS 40:4 | 40 | 4 |
| PS 40:6 | 40 | 6 |
| SM 30:1 | 30 | 1 |
| SM 32:1 | 32 | 1 |
| SM 34:1 | 34 | 1 |
| SM 34:2 | 34 | 2 |
| SM 35:1 | 35 | 1 |
| SM 35:2 | 35 | 2 |
| SM 39:1 | 39 | 1 |

| | | |
|---|---|---|
| SM 40:1 | 40 | 1 |
| SM 40:1 | 40 | 1 |
| SM 41:1 | 41 | 1 |
| SM 41:4 | 41 | 4 |
| SM 41:5 | 41 | 5 |
| SM 42:2 | 42 | 2 |
| TG 40:0 | 40 | 0 |
| TG 42:0 | 42 | 0 |
| TG 46:0 | 46 | 0 |
| TG 48:0 | 48 | 0 |
| TG 48:1 | 48 | 1 |
| TG 49:0 | 49 | 0 |
| TG 49:1 | 49 | 1 |
| TG 50:0 | 50 | 0 |
| TG 50:1 | 50 | 1 |
| TG 50:4 | 50 | 4 |
| TG 51:1 | 51 | 1 |
| TG 52:0 | 52 | 0 |
| TG 52:1 | 52 | 1 |
| TG 52:5 | 52 | 5 |
| TG 53:0 | 53 | 0 |
| TG 53:2 | 53 | 2 |
| TG 54:0 | 54 | 0 |
| TG 54:1 | 54 | 1 |
| TG 54:2 | 54 | 2 |
| TG 54:7 | 54 | 7 |
| TG 54:8 | 54 | 8 |
| TG 55:3 | 55 | 3 |
| TG 56:0 | 56 | 0 |
| TG 56:1 | 56 | 1 |
| TG 56:2 | 56 | 2 |
| TG 56:3 | 56 | 3 |
| TG 56:4 | 56 | 4 |
| TG 56:5 | 56 | 5 |
| TG 58:1 | 58 | 1 |
| TG 58:2 | 58 | 2 |
| TG 58:3 | 58 | 3 |
| TG 58:4 | 58 | 4 |
| TG 58:5 | 58 | 5 |
| TG 58:6 | 58 | 6 |
| TG 60:1 | 60 | 1 |
| TG 60:2 | 60 | 2 |
| TG 60:3 | 60 | 3 |

| | | |
|---|---|---|
| TG 60:4 | 60 | 4 |
| TG 60:5 | 60 | 5 |
| TG 60:6 | 60 | 6 |
| TG 60:7 | 60 | 7 |
| TG 62:8 | 62 | 8 |

**Supplementary Table 2. List of differential lipids (FDR<0.25) in AA PCa vs matched benign adjacent tissues.** List of detected differential lipids between AA PCa and matched benign adjacent tissue arranged by lipid name, class, length, and bond number. PE: Phosphatidyl ethanolamine, PC: Phosphatidyl choline, PI: Phosphatidyl inositol, PG: Phosphatidyl Glycerol, PS: Phosphatidyl Serine, L-PE: Lyso-Phosphatidyl Ethanolamine, TG: Triglycerides, SM: Sphingomyelin, P-PE: Plasmenyl-Phosphatidyl Ethanolamine, CE: Cholesteryl Esters, DG: Diglycerides, CL: Cardiolipins, L-PC: Lyso-Phosphatidyl Choline. The fatty acid chain length and the number of bonds (degree of saturation) are indicated.

| **Lipid Name** | **Length of chain** | **Bond Number** |
|---|---|---|
| CE 16:1; [M+NH4]+@10.16 | 16 | 1 |
| CE 18:0; [M+NH4]+@10.84 | 18 | 0 |
| CE 18:1; [M+NH4]+@10.51 | 18 | 1 |
| CE 18:3; [M+NH4]+@10.03 | 18 | 3 |
| CE 19:0; [M+NH4]+@11.00 | 19 | 0 |
| CE 20:0; [M+NH4]+@11.12 | 20 | 0 |
| CE 20:1; [M+NH4]+@10.85 | 20 | 1 |
| CE 20:3; [M+NH4]+@10.30 | 20 | 3 |
| CE 20:5; [M+NH4]+@9.89 | 20 | 5 |
| CE 22:1; [M+NH4]+@11.23 | 22 | 1 |
| CE 22:2; [M+NH4]+@10.87 | 22 | 2 |
| CE 22:4; [M+NH4]+@10.40 | 22 | 4 |
| CE 22:5; [M+NH4]+@10.22 | 22 | 5 |
| CE 22:6; [M+NH4]+@10.00 | 22 | 6 |
| CE 24:1; [M+NH4]+@11.39 | 24 | 1 |
| DG 32:0; [M+NH4]+@7.12 | 32 | 0 |
| DG 33:0; [M+NH4]+@7.45 | 33 | 0 |
| DG 34:0; [M+NH4]+@7.66 | 34 | 0 |
| DG 35:0; [M+NH4]+@7.89 | 35 | 0 |
| DG 35:1; [M+NH4]+@7.42 | 35 | 1 |
| DG 35:2; [M+NH4]+@6.99 | 35 | 2 |
| DG 36:0; [M+NH4]+@8.13 | 36 | 0 |
| DG 37:1; [M+NH4]+@8.03 | 37 | 1 |
| DG 37:2; [M+NH4]+@7.64 | 37 | 2 |
| DG 37:4; [M+NH4]+@7.03 | 37 | 4 |
| DG 38:1; [M+NH4]+@8.17 | 38 | 1 |

| | | |
|---|---|---|
| DG 38:2; [M+NH4]+@7.76 | 38 | 2 |
| DG 38:4; [M+NH4]+@7.17 | 38 | 4 |
| DG 38:5; [M+NH4]+@6.73 | 38 | 5 |
| DG 40:2; [M+NH4]+@8.18 | 40 | 2 |
| DG 40:4; [M+NH4]+@7.63 | 40 | 4 |
| DG 40:5; [M+NH4]+@7.30 | 40 | 5 |
| lysoPC 17:0; [M+H]+@2.40 | 17 | 0 |
| lysoPE 16:1; [M-H]-@1.19 | 16 | 1 |
| lysoPE 17:0; [M-H]-@2.04 | 17 | 0 |
| lysoPE 18:0; [M-H]-@2.40 | 18 | 0 |
| lysoPE 18:1; [M-H]-@1.76 | 18 | 1 |
| lysoPE 18:2; [M-H]-@1.22 | 18 | 2 |
| lysoPE 19:0; [M-H]-@2.98 | 19 | 0 |
| lysoPE 20:0; [M-H]-@3.44 | 20 | 0 |
| lysoPE 22:1; [M+H]+@2.55 | 22 | 1 |
| PC 28:0; [M+H]+@4.73 | 28 | 0 |
| PC 31:0; [M-Ac-H]-@6.28 | 31 | 0 |
| PC 32:0; [M+Na]+@6.4075 | 32 | 0 |
| PC 33:0; [M-Ac-H]-@6.91 | 33 | 0 |
| PC 35:1; [M-Ac-H]-@7.07 | 35 | 1 |
| PC 36:0; [M-Ac-H]-@7.85 | 36 | 0 |
| PC 36:3; [M+H]+@7.06 | 36 | 3 |
| PC 36:4; [M+H]+@5.835 | 36 | 4 |
| PC 37:6; [M-Ac-H]-@5.83 | 37 | 6 |
| PC 38:1; [M-Ac-H]-@7.88 | 38 | 1 |
| PC 38:6; [M-Ac-H]-@6.00 | 38 | 6 |
| PC 40:6; [M+Na]+@6.3825 | 40 | 6 |
| PC 40:7; [M+H]+@9.10 | 40 | 7 |
| PC 40:8; [M-Ac-H]-@5.80 | 40 | 8 |
| PC 42:10; [M-Ac-H]-@5.72 | 42 | 10 |
| PC 42:6; [M-Ac-H]-@7.36 | 42 | 6 |
| PC 42:9; [M+H]+@6.13 | 42 | 9 |
| PE 30:0; [M+H]+@5.40 | 30 | 0 |
| PE 32:2; [M+H]+@4.98 | 32 | 2 |
| PE 34:0; [M+Na]+@6.695 | 34 | 0 |
| PE 36:1; [M+Na]+@6.77 | 36 | 1 |
| PE 36:3; [M-H]-@6.55 | 36 | 3 |
| PE 36:4; [M-H]-@7.66 | 36 | 4 |
| PE 37:2; [M-H]-@7.28 | 37 | 2 |
| PE 37:4; [M-H]-@6.64 | 37 | 4 |
| PE 38:4; [M-H]-@7.86 | 38 | 4 |
| PE 39:7; [M-H]-@6.10 | 39 | 7 |
| PE 40:1; [M+H]+@7.88 | 40 | 1 |

| | | |
|---|---|---|
| PE 40:4; [M+H]+@6.94 | 40 | 4 |
| PE 40:8; [M-H]-@6.00 | 40 | 8 |
| PE 41:4; [M-H]-@7.66 | 41 | 4 |
| PE 41:6; [M-H]-@7.21 | 41 | 6 |
| PE 42:4; [M-H]-@7.94 | 42 | 4 |
| PE 42:6; [M-H]-@7.46 | 42 | 6 |
| PE 42:7; [M-H]-@6.96 | 42 | 7 |
| PE 44:4; [M-H]-@8.53 | 44 | 4 |
| PE 44:5; [M-H]-@8.09 | 44 | 5 |
| PG 34:0; [M-H]-@6.295 | 34 | 0 |
| PG 36:1; [M-H]-@6.45 | 36 | 1 |
| PG 36:2; [M-H]-@5.85 | 36 | 2 |
| PG 36:3; [M-H]-@5.37 | 36 | 3 |
| PG 38:3; [M-H]-@5.81 | 38 | 3 |
| PG 38:4; [M-H]-@5.44 | 38 | 4 |
| PG 38:4; [M-H]-@6.09 | 38 | 4 |
| PG 38:5; [M-H]-@5.15 | 38 | 5 |
| PG 42:6; [M-H]-@6.07 | 42 | 6 |
| PI 36:2; [M-H]-@5.90 | 36 | 2 |
| PI 38:3; [M-H]-@6.13 | 38 | 3 |
| PI 39:4; [M-H]-@6.23 | 39 | 4 |
| plasmenyl-PE 32:0; [M+H]+@6.41 | 32 | 0 |
| plasmenyl-PE 36:1; [M+Na]+@7.075 | 36 | 1 |
| plasmenyl-PE 36:4; [M+H]+@5.85 | 36 | 4 |
| plasmenyl-PE 36:4; [M-H]-@6.67 | 36 | 4 |
| plasmenyl-PE 36:5; [M+H]+@4.04 | 36 | 5 |
| plasmenyl-PE 36:5; [M+H]+@5.50 | 36 | 5 |
| plasmenyl-PE 36:5; [M-H]-@6.35 | 36 | 5 |
| plasmenyl-PE 38:4; [M+Na]+@6.875 | 38 | 4 |
| plasmenyl-PE 38:5; [M+H]+@5.91 | 38 | 5 |
| plasmenyl-PE 38:5; [M-H]-@6.76 | 38 | 5 |
| plasmenyl-PE 38:6; [M+H]+@5.77 | 38 | 6 |
| plasmenyl-PE 38:6; [M-H]-@6.60 | 38 | 6 |
| plasmenyl-PE 40:4; [M+Na]+@7.3625 | 40 | 4 |
| plasmenyl-PE 40:5; [M+H]+@6.63 | 40 | 5 |
| plasmenyl-PE 40:5; [M-H]-@7.47 | 40 | 5 |
| plasmenyl-PE 42:4; [M-H]-@8.09 | 42 | 4 |
| PS 36:2; [M+Na]+@4.94 | 36 | 2 |
| PS 36:4; [M-H]-@5.32 | 36 | 4 |
| PS 38:4; [M+Na]+@4.92 | 38 | 4 |

| | | |
|---|---|---|
| PS 38:5; [M-H]-@5.47 | 38 | 5 |
| PS 38:6; [M-H]-@5.22 | 38 | 6 |
| PS 38:7; [M+H]+@4.38 | 38 | 7 |
| PS 40:4; [M-H]-@6.40 | 40 | 4 |
| PS 40:6; [M+Na]+@4.78 | 40 | 6 |
| SM 30:1; [M]+@3.51 | 30 | 1 |
| SM 35:1; [M+Na]+@8.43 | 35 | 1 |
| SM 35:2; [M]+@5.07 | 35 | 2 |
| SM 39:1; [M]+@7.17 | 39 | 1 |
| SM 40:2; [M]+@7.04 | 40 | 2 |
| SM 41:1; [M+Na]+@7.895 | 41 | 1 |
| SM 41:4; [M]+@6.18 | 41 | 4 |
| SM 41:5; [M]+@6.10 | 41 | 5 |
| SM 42:0; [M+Na]+@9.78 | 42 | 0 |
| SM 42:2; [M]+@9.01 | 42 | 2 |
| TG 40:0; [M+NH4]+@8.83 | 40 | 0 |
| TG 42:0; [M+NH4]+@9.19 | 42 | 0 |
| TG 46:0; [M+NH4]+@9.785 | 46 | 0 |
| TG 48:0; [M+NH4]+@10.07 | 48 | 0 |
| TG 48:1; [M+NH4]+@9.815 | 48 | 1 |
| TG 49:0; [M+NH4]+@10.15 | 49 | 0 |
| TG 49:1; [M+NH4]+@9.96 | 49 | 1 |
| TG 50:0; [M+NH4]+@10.33 | 50 | 0 |
| TG 50:1; [M+NH4]+@10.11 | 50 | 1 |
| TG 50:5; [M+NH4]+@9.10 | 50 | 5 |
| TG 51:1; [M+NH4]+@10.225 | 51 | 1 |
| TG 52:0; [M+NH4]+@10.6 | 52 | 0 |
| TG 52:1; [M+NH4]+@10.325 | 52 | 1 |
| TG 53:0; [M+NH4]+@10.685 | 53 | 0 |
| TG 53:1; [M+NH4]+@10.505 | 53 | 1 |
| TG 53:2; [M+NH4]+@10.25 | 53 | 2 |
| TG 54:1; [M+NH4]+@10.6 | 54 | 1 |
| TG 54:2; [M+NH4]+@10.365 | 54 | 2 |
| TG 54:7; [M+NH4]+@9.285 | 54 | 7 |
| TG 55:1; [M+NH4]+@10.77 | 55 | 1 |
| TG 55:2; [M+NH4]+@10.47 | 55 | 2 |
| TG 55:3; [M+NH4]+@10.31 | 55 | 3 |
| TG 55:6; [M+NH4]+@9.77 | 55 | 6 |
| TG 56:0; [M+NH4]+@11.035 | 56 | 0 |
| TG 56:1; [M+NH4]+@10.82 | 56 | 1 |
| TG 56:2; [M+NH4]+@10.62 | 56 | 2 |
| TG 56:3; [M+NH4]+@10.4 | 56 | 3 |
| TG 56:4; [M+NH4]+@10.235 | 56 | 4 |

| | | |
|---|---|---|
| TG 58:1; [M+NH4]+@11.06 | 58 | 1 |
| TG 58:2; [M+NH4]+@10.82 | 58 | 2 |
| TG 58:3; [M+NH4]+@10.63 | 58 | 3 |
| TG 58:4; [M+NH4]+@10.44 | 58 | 4 |
| TG 58:5; [M+NH4]+@10.295 | 58 | 5 |
| TG 58:6; [M+NH4]+@10.11 | 58 | 6 |
| TG 60:1; [M+NH4]+@11.26 | 60 | 1 |
| TG 60:2; [M+NH4]+@11.06 | 60 | 2 |
| TG 60:3; [M+NH4]+@10.85 | 60 | 3 |
| TG 60:4; [M+NH4]+@10.67 | 60 | 4 |
| TG 60:5; [M+NH4]+@10.56 | 60 | 5 |
| TG 60:6; [M+NH4]+@10.34 | 60 | 6 |
| TG 60:7; [M+NH4]+@10.16 | 60 | 7 |

**Supplementary Table 3. List of differential lipids (FDR<0.25) comparing AA vs EA PCa tumors.** List of detected differential lipids between AA vs EA PCa tissues arranged by lipid name, length, and bond number. PE: Phosphatidyl ethanolamine, PC: Phosphatidyl choline, PI: Phosphatidyl inositol, PG: Phosphatidyl Glycerol, PS: Phosphatidyl Serine, L-PE: Lyso-Phosphatidyl Ethanolamine, TG: Triglycerides, SM: Sphingomyelin, P-PE: Plasmenyl-Phosphatidyl Ethanolamine, CE: Cholesteryl Esters, DG: Diglycerides, CL: Cardiolipins, L-PC: Lyso-Phosphatidyl Choline. The fatty acid chain length and the number of bonds (degree of saturation) are indicated.

| **Lipid Name** | **Chain Length** | **Bond Number** |
|---|---|---|
| CE 16:1 [M+NH4]+@10.16 | 16 | 1 |
| CE 18:0 [M+NH4]+@10.84 | 18 | 0 |
| CE 18:1 [M+NH4]+@10.51 | 18 | 1 |
| CE 19:0 [M+NH4]+@11.00 | 19 | 0 |
| CE 20:0 [M+NH4]+@11.12 | 20 | 0 |
| CE 20:1 [M+NH4]+@10.85 | 20 | 1 |
| CE 20:5 [M+NH4]+@9.89 | 20 | 5 |
| CE 22:1 [M+NH4]+@11.23 | 22 | 1 |
| CE 22:2 [M+NH4]+@10.87 | 22 | 2 |
| CE 22:5 [M+NH4]+@10.22 | 22 | 5 |
| CE 22:6 [M+NH4]+@10.00 | 22 | 6 |
| PA 34:0 [M-H]-@6.71 | 34 | 0 |
| PA 37:4 [M-H]-@5.71 | 37 | 4 |
| PC 32:0 [M+Na]+@6.4075 | 32 | 0 |
| PC 42:10 [M-Ac-H]-@5.72 | 42 | 10 |
| PG 36:2 [M-H]-@5.85 | 36 | 2 |
| PI 36:2 [M-H]-@5.90 | 36 | 2 |
| PI 36:4 [M-H]-@5.25 | 36 | 4 |

| | | |
|---|---|---|
| PI 38:3 [M-H]-@6.13 | 38 | 3 |
| PI 38:4 [M-H]-@5.91 | 38 | 4 |
| PI 39:4 [M-H]-@6.23 | 39 | 4 |
| PI 40:4 [M-H]-@6.50 | 40 | 4 |
| plasmenyl-PC 20:0 [M+Na]+@2.02 | 20 | 0 |
| plasmenyl-PE 32:0 [M+H]+@6.41 | 32 | 0 |
| SM 41:4 [M]+@6.18 | 41 | 4 |
| SM 43:1 [M]+@8.60 | 43 | 1 |
| SM 43:1 [M+Na]+@8.74 | 43 | 1 |
| SM 44:1 [M]+@8.94 | 44 | 1 |
| TG 55:1 [M+NH4]+@10.77 | 55 | 1 |
| TG 60:5 [M+NH4]+@10.56 | 60 | 5 |
