## Supplementary Figures for "Lipid Alterations in African American Prostate Cancer"

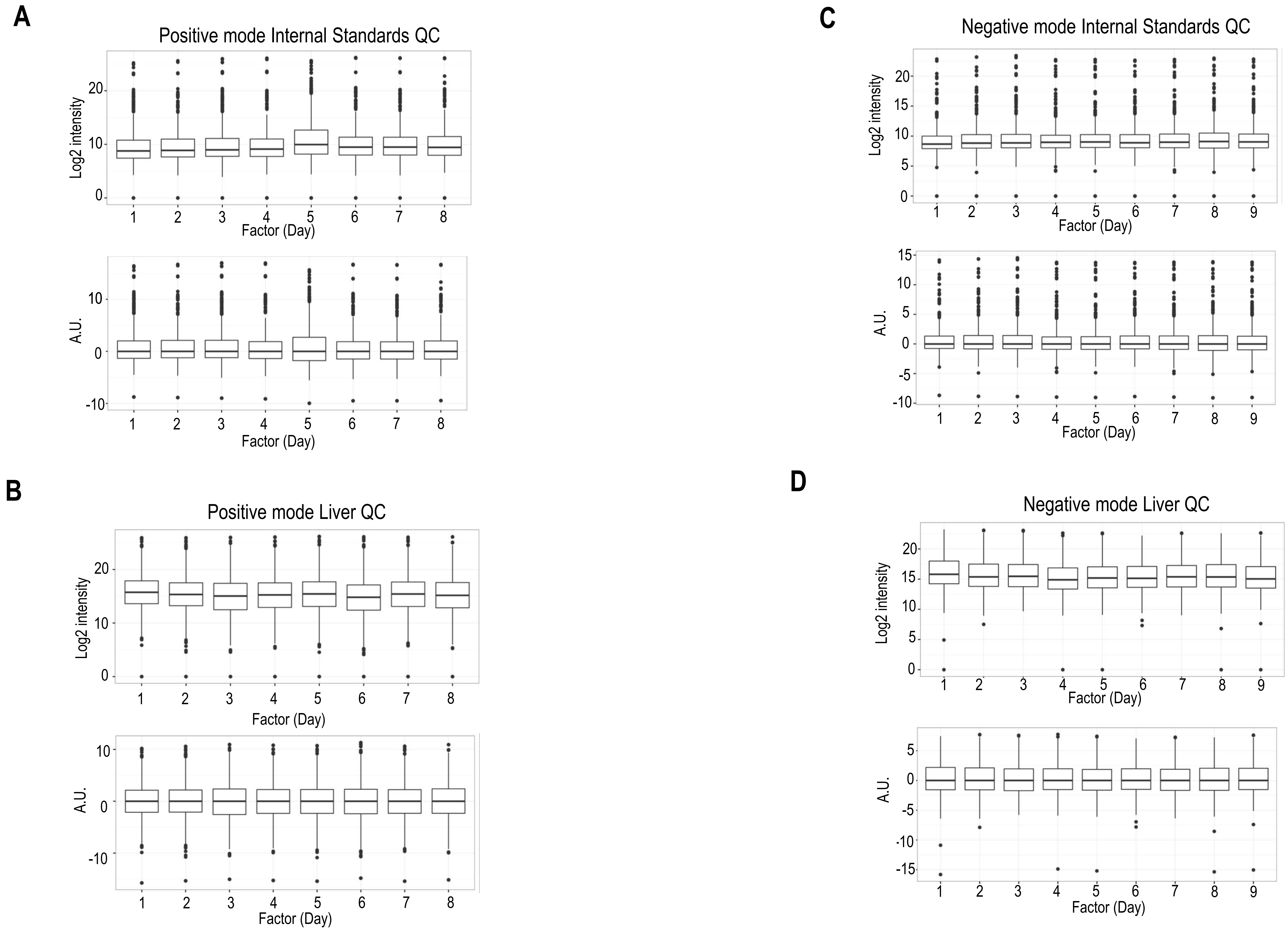


**Supplementary Figure 1.** Quality control data for the lipidomics profiling mass spectrometry platform. **(A)** and **(B)** show distribution of internal standards for positive and negative ionization modes. Lower panel shows the raw data in arbitrary units (AU) across the nine days of analysis. The upper panel shows the normalized data in log2 scale. **(C)** and **(D)** similar to (A) and (B), but for liver pools. Three liver pool samples were run per day across eight days.

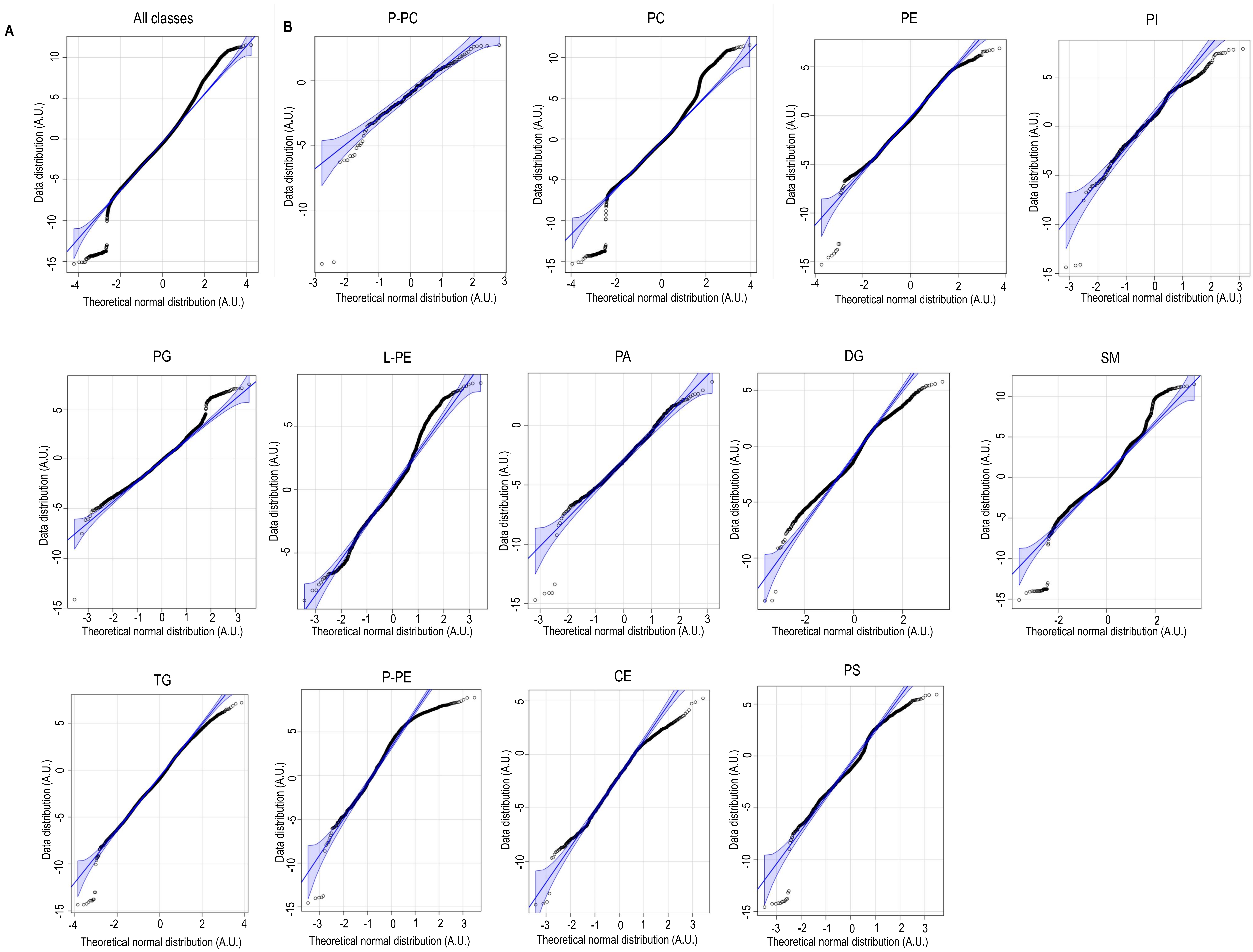


**Supplementary Figure 2.** Quantile-quantile plots for **(A)** All lipid classes and **(B)** individual lipid classes. P-PC: Plasmenyl-Phosphatidyl Choline, PE: Phosphatidyl ethanolamine, PC: Phosphatidyl choline, PI: Phosphatidyl inositol, PG: Phosphatidyl Glycerol, PS: Phosphatidyl Serine, L-PE: Lyso-Phosphatidyl Ethanolamine, TG: Triglycerides, SM: Sphingomyelin, P-PE: Plasmenyl-Phosphatidyl Ethanolamine, CE: Cholesteryl Esters, DG: Diglycerides, L-PC: Lyso-Phosphatidyl Choline, PA: Phosphatidic Acid.

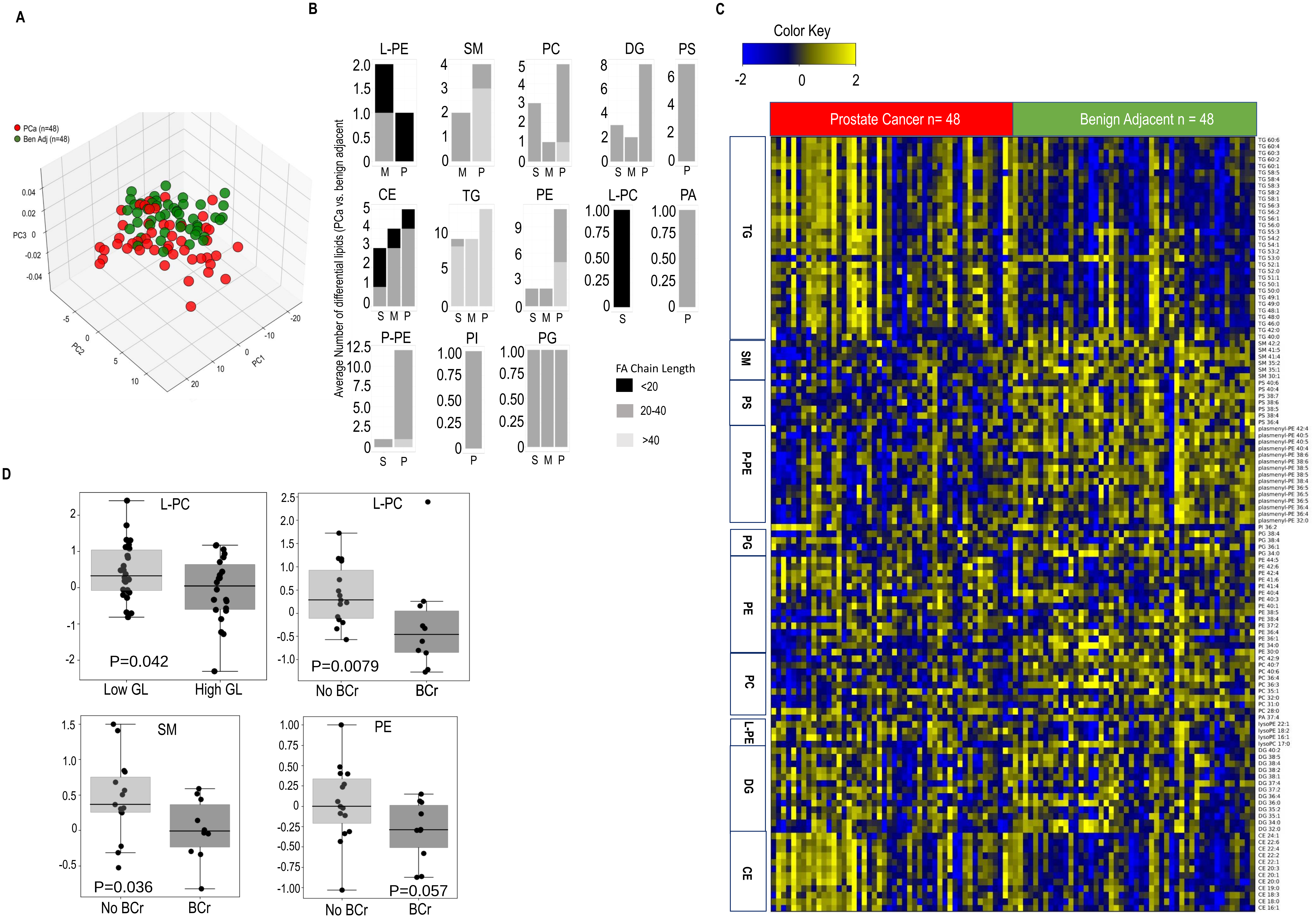


**Supplemental Figure 3. Altered lipidome in Prostate Cancer (PCa) vs matched benign adjacent tissue. (A)** PCA plot using lipid profiles in 48 paired patient-derived PCa and benign adjacent tissues. **(B)** Number of altered lipids within each class stratified by fatty acid chain length (see legend) and degree of saturation. S: Saturated, M: Mono-unsaturated, P: Poly-unsaturated (≥ 2 double bonds). **(C)** Heat map showing significantly altered lipids in PCa vs matched benign adjacent tissues. Shades of Yellow and Blue represent up and down regulated lipids, respectively (see color key). Lipids are arranged by classes. PE: Phosphatidyl ethanolamine, PC: Phosphatidyl choline, PI: Phosphatidyl inositol, PG: Phosphatidyl Glycerol, PS: Phosphatidyl Serine, L-PE: Lyso-Phosphatidyl Ethanolamine, TG: Triglycerides, SM: Sphingomyelin, P-PE: Plasmenyl-Phosphatidyl Ethanolamine, CE: Cholesteryl Esters, DG: Diglycerides, L-PC: Lyso-Phosphatidyl Choline. **(D)** Levels of L-PC (p=0.042) are significantly down-regulated in high Gleason grade (High GL, n=26) compared to Low Gleason grade (Low GL, n=22) tumors. Along similar lines, lower levels of L-PC (p=0.0079) SM (p=0.036) and PE (p=0.057) are associated with biochemical recurrence (BCr within 5 years post-prostatectomy) in PCa patients. No BCr (n=15); BCr (n=10). For panels B and C, paired t-test followed by Benjamini Hochberg (BH) False Discovery Rate (FDR<0.1) correction was used to compute differential analysis. For panel D, Mann-Whitney test with BH FDR<0.25 (except for BCr analysis where FDR<0.35) was used to compute statistical significance.

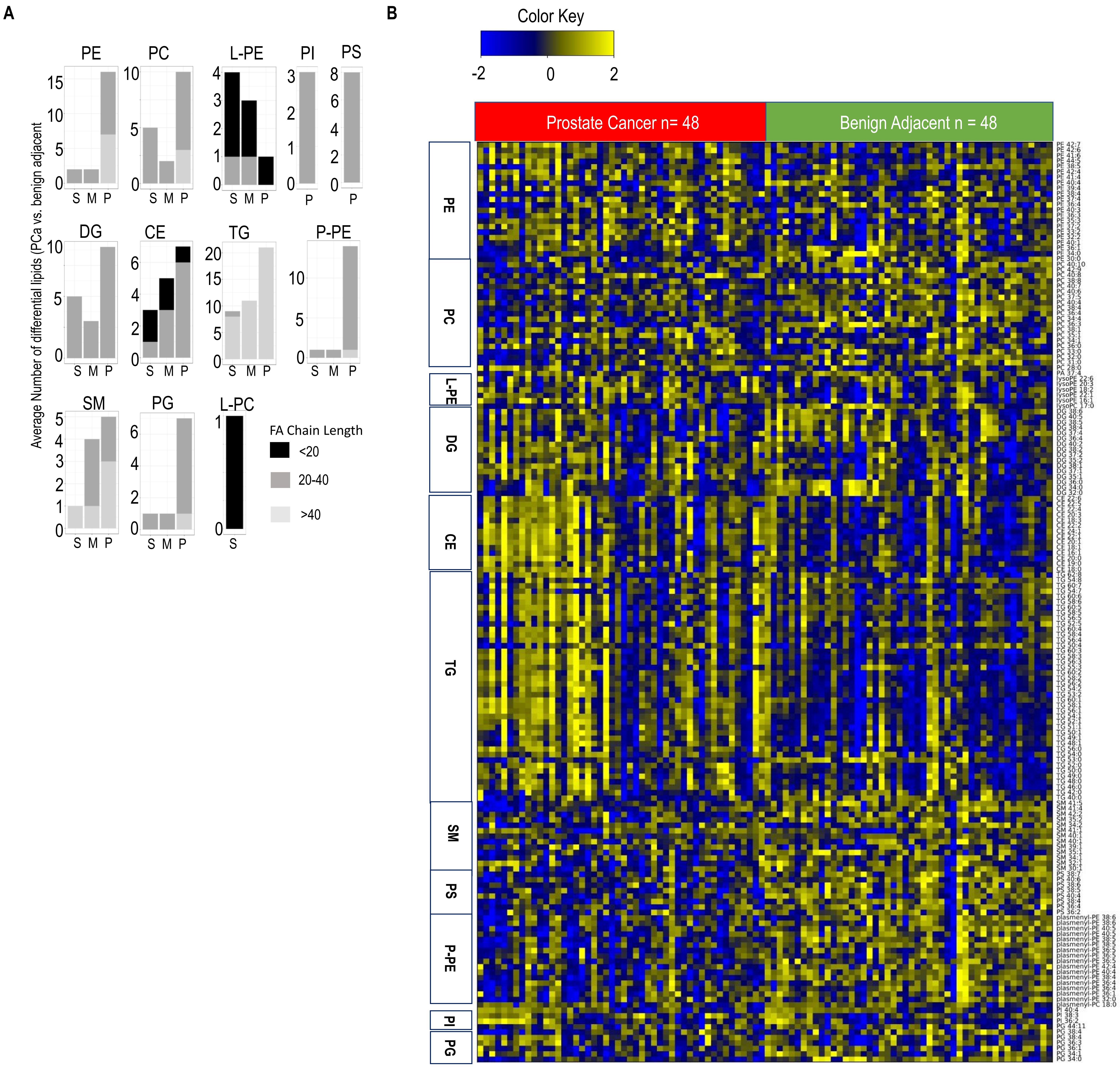


**Supplemental Figure 4. Altered lipidome in Prostate Cancer (PCa) vs matched benign adjacent tissue (FDR<0.25). (A)** Number of altered lipids within each class stratified by fatty acid chain length (see legend) and degree of saturation. S: Saturated, M: Mono-unsaturated, P: Poly-unsaturated (≥ 2 double bonds). **(B)** Heat map showing significantly altered lipids in PCa vs matched benign adjacent tissues. Shades of Yellow and Blue represent up and down regulated lipids, respectively (see color key). Lipids are arranged by classes. PE: Phosphatidyl ethanolamine, PC: Phosphatidyl choline, PI: Phosphatidyl inositol, PG: Phosphatidyl Glycerol, PS: Phosphatidyl Serine, L-PE: Lyso-Phosphatidyl Ethanolamine, TG: Triglycerides, SM: Sphingomyelin, P-PE: Plasmenyl-Phosphatidyl Ethanolamine, CE: Cholesteryl Esters, DG: Diglycerides, L-PC: Lyso-Phosphatidyl Choline. For panels A and B, paired t-test followed by Benjamini Hochberg (BH) False Discovery Rate (FDR<0.25) correction was used to compute differential analysis.

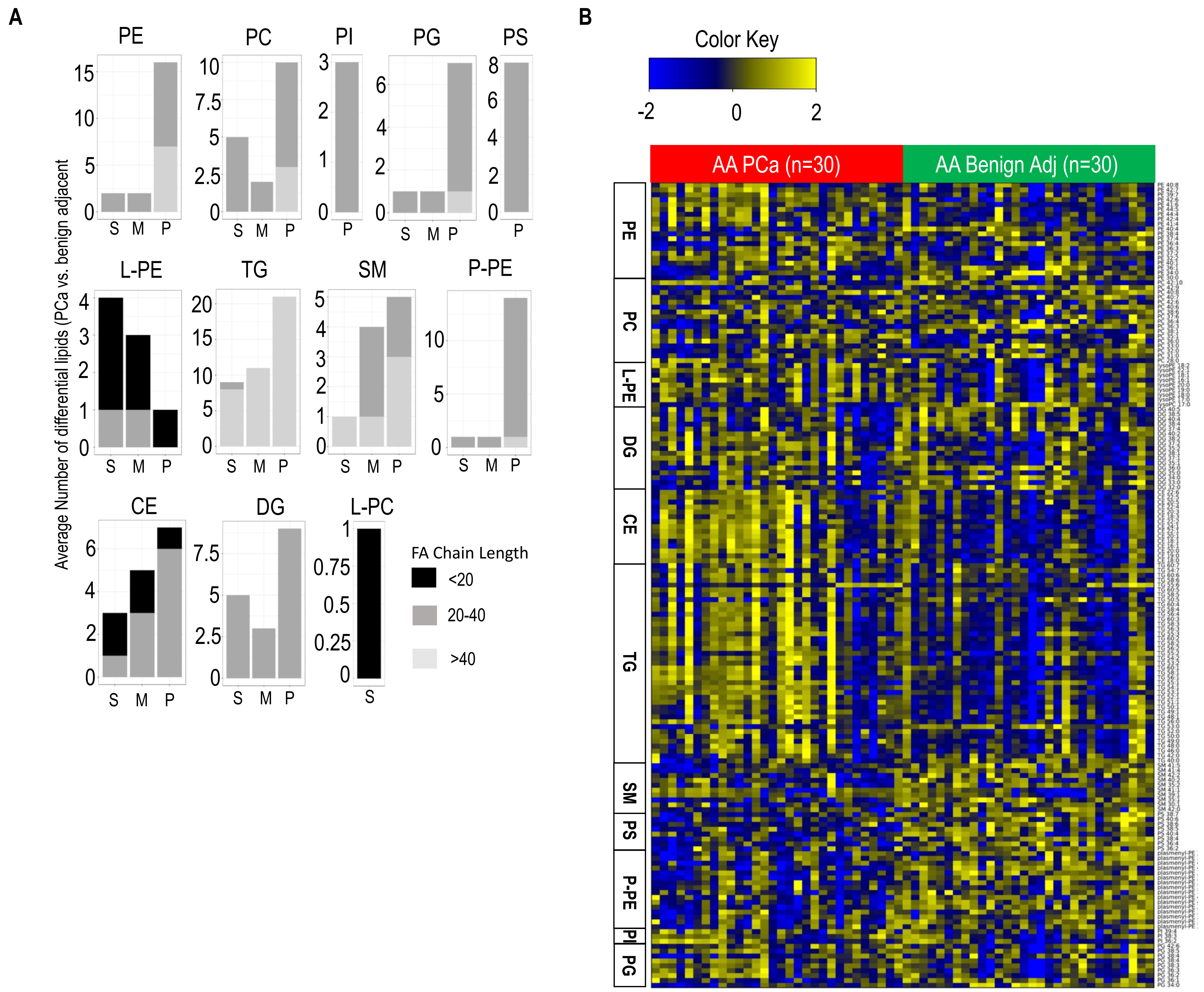


**Supplemental Figure 5. Altered lipidome in AA PCa vs matched benign patient prostate tissue (FDR<0.25). (A)** Number of altered lipids within each class stratified by fatty acid chain length (see legend) and degree of saturation. S: Saturated, M: Mono-unsaturated, P: Poly-unsaturated (≥ 2 double bonds). **(B)** Heat map showing significantly altered lipids in AA PCa vs matched benign patient prostate tissues. Shades of Yellow and Blue represent up and down regulated lipids, respectively (see color key). Lipids are arranged by classes. PE: Phosphatidyl ethanolamine, PC: Phosphatidyl choline, PI: Phosphatidyl inositol, PG: Phosphatidyl Glycerol, PS: Phosphatidyl Serine, L-PE: Lyso Phosphatidyl Ethanolamine, TG: Triglycerides, SM: Sphingomyelin, P-PE: Plasmenyl Phosphatidyl Ethanolamine, CE: Cholesteryl Esters, DG: Diglycerides, L-PC: Lyso phosphatidyl choline. For panels A and B, paired t-test followed by Benjamini Hochberg (BH) False Discovery Rate (FDR<0.25) correction was used to compute differential analysis.

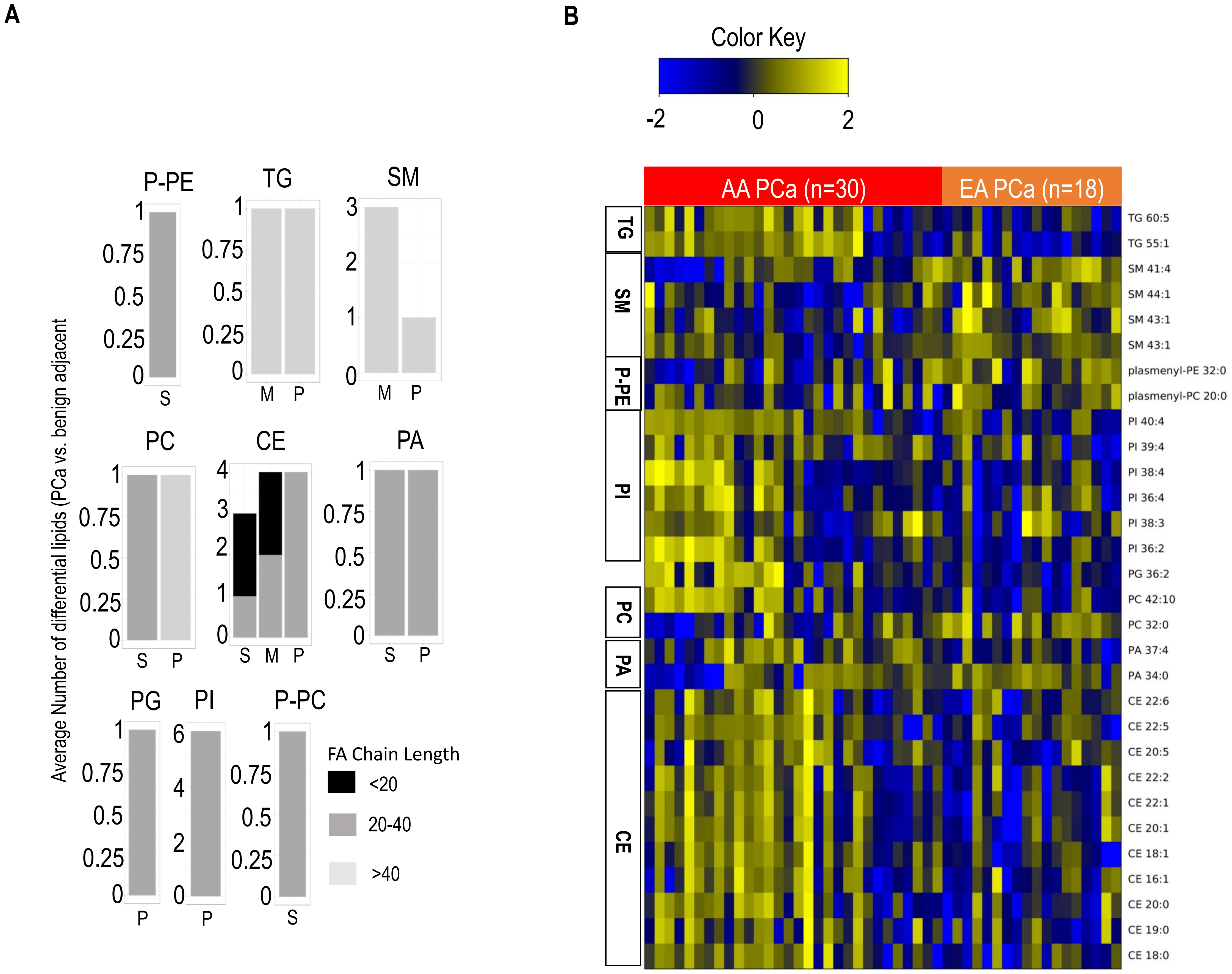


**Supplemental Figure 6. Altered lipidome in AA PCa vs EA PCa patient prostate cancer tissue (FDR<0.25). (A)** Number of altered lipids within each class stratified by fatty acid chain length (see legend) and degree of saturation. S: Saturated, M: Mono-unsaturated, P: Poly-unsaturated (≥ 2 double bonds). **(B)** Heat map showing significantly altered lipids in AA PCa vs EA PCa tissues. Shades of Yellow and Blue represent up and down regulated lipids, respectively (see color key). Lipids are arranged by classes. P-PC: Plasmenyl Phosphatidyl choline, P-PE: Plasmenyl Phosphatidyl Ethanolamine, TG: Triglycerides, SM: Sphingomyelin, PI: Phosphatidyl inositol, PG: Phosphatidyl Glycerol, PC: Phosphatidyl choline, CE: Cholesteryl Esters, PA: Phosphatidic Acid. For panels B and C, paired t-test coupled to Benjamin Hochberg False Discovery Rate (FDR<0.25) correction was used to compute differential analysis.
